# Molecular characterization of CHAD domains as inorganic polyphosphate binding modules

**DOI:** 10.1101/567040

**Authors:** Laura Lorenzo-Orts, Ulrich Hohmann, Jinsheng Zhu, Michael Hothorn

**Affiliations:** Structural Plant Biology Laboratory, Department of Botany and Plant Biology, Faculty of Sciences, University of Geneva, 1211 Geneva 4, Switzerland

## Abstract

Inorganic polyphosphates (polyPs) are long polymers of orthophosphate units (P_i_), linked by energy-rich phosphoanhydride bonds. Conserved histidine α-helical (CHAD) domains of unknown biochemical function are often located at the C-terminus of polyP-metabolizing triphosphate tunnel metalloenzymes (TTMs), or can be found as stand-alone proteins in bacterial operons harboring polyP kinases or phosphatases. Here we report that bacterial, archaeal and eukaryotic CHAD domains are specific polyP binding modules. Crystal structures reveal that CHAD domains are formed by two four-helix bundles, giving rise to a central cavity surrounded by two conserved basic surface patches. Different CHAD domains bind polyPs with dissociation constants ranging from the nano-to mid-micromolar range, but not DNA or other P_i_-containing ligands. A 2.1 Å CHAD - polyP complex structure reveals the phosphate polymer binding across a central pore and along the two basic patches. Mutational analysis of CHAD – polyP interface residues validates the complex structure and reveals that CHAD domains evolved to bind long-chain polyPs. The presence of a CHAD domain in the polyPase ygiF enhances its enzymatic activity. In plants, CHAD domains bind polyP *in vivo* and localize to the nucleus and nucleolus, suggesting that plants harbor polyP stores in these compartments. We propose that CHAD domains may be used to engineer the properties of polyP-metabolizing enzymes and to specifically localize polyP stores in eukaryotic cells and tissues.

**Significance:** A domain of unknown function termed CHAD, present in all kingdoms of life, is characterized as a specific inorganic polyphosphate binding domain. The small size of the domain and its high specificity for inorganic polyphosphates suggest that it could be used as a tool to locate inorganic polyphosphate stores in pro- and eukaryotic cells and tissues.

## Introduction

Inorganic polyphosphates (polyPs) form an important phosphate (P_i_) and energy store in pro- and eukaryotic cells (1, 2). In bacteria, polyPs can form granules in the nucleoid region, regulate the cell cycle (3), form cation-selective membrane channels (4), control cell motility (5), mediate cellular stress responses, for instance by preventing protein aggregation (6). In eukaryotes, polyPs have thus far been found in vacuoles or specialized acidocalcisomes (7) and form an important store for P_i_ (8–11) and divalent metal ions (12, 13). At the physiological level, polyPs are involved in cell cycle control (14), in cell death responses (15), coagulation (16), skeletal mineralization (17) and in the post-translational modification of proteins (18).

PolyP-metabolizing enzymes have been well-characterized in bacteria and lower eukaryotes. In bacteria polyP may be synthesized from ATP by the polyphosphate kinase 1 (PPK1) (19, 20) or from ATP/GTP by PPK2 (21–24). In lower eukaryotes such as fungi, protozoa and algae, polyP is generated from ATP by the membrane-integral VTC complex (9, 25–27). No polyphosphate kinase has been reported from higher eukaryotes thus far, despite the presence of polyPs in these organisms (28). Exopolyphosphatase PPX1 (29), and the triphosphate tunnel metalloenzyme (TTM) ygiF (30) are polyP degrading enzymes in bacteria. Eukaryotic polyphosphatases include the yeast exopolyphosphatase 1 (PPX1) (31), the endopolyphosphatases PPN1 (32) and PPN2 (33), the Ddp1-type Nudix hydrolases (34), human H-prune (35) and the plant tripolyphosphatase TTM3 (36, 30).

To date no polyP binding domain has been identified, although an engineered polyP binding domain from EcPPX1 has used to immunolocalize polyPs in eukaryotic cells and tissues (37). We have previously identified a small, helical domain at the C-terminus of the bacterial short-chain polyphosphatase ygiF (30, 38). This domain of unknown function has been annotated as CHAD (conserved histidine α-helical domain, PFAM PF05235) (39). Many CHAD domain-containing proteins harbor a N-terminal CYTH/TTM domain, while stand-alone CHAD proteins often are part of operons expressing polyP metabolic enzymes (39). Recently, it was found that CHAD domain-containing proteins specifically localize to polyP granules in the bacterium *Ralstonia eutropha* (40). In this study, we combine structural biology and quantitative biochemistry to define CHAD domains as polyP binding modules.

## Results

We located CHAD domains in the different kingdoms of life. According to Interpro (https://www.ebi.ac.uk/interpro), ∼ 99 % of the annotated CHAD proteins correspond to bacteria, while only ∼ 1 % (129 proteins) and 0.1 % (10 proteins) belong to archaea and eukaryota, respectively (Fig. 1*A*). We selected CHAD domain-containing proteins belonging to the three kingdoms of life: archaea (*Sulfolobus solfataricus;* termed SsCHAD hereafter), bacteria (*Chlorobium tepidum;* CtCHAD) and eukaryota (*Ricinus communis* or castor bean; RcCHAD) (Fig 1*A*) (*SI Appendix* Fig. S1). Several of these CHAD proteins form part of gene clusters encoding polyP metabolizing enzymes, with the exception of RcCHAD (Fig. 1*B*).

**Figure 1.**
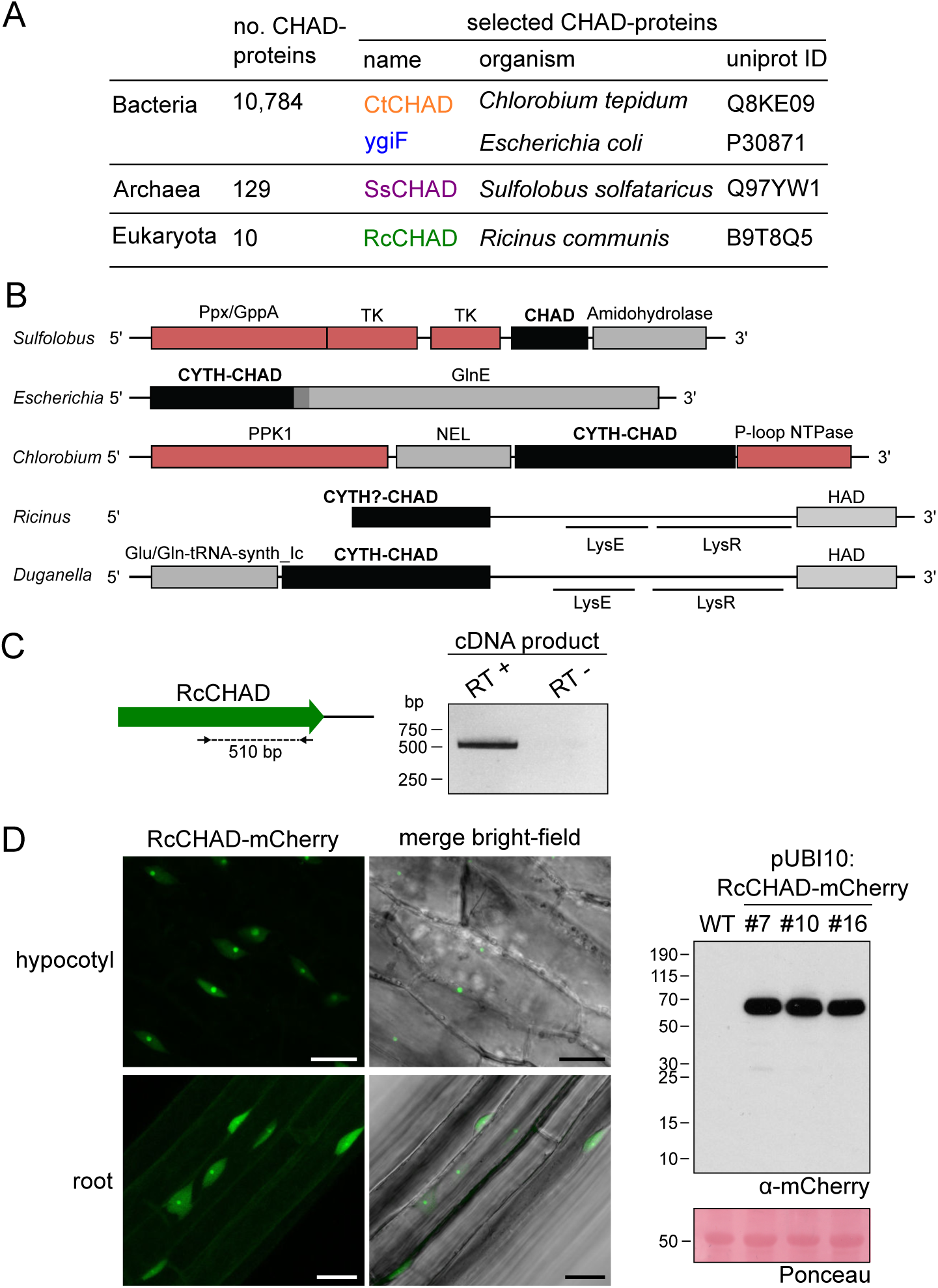
CHAD domain-containing proteins are present in all kingdoms of life. **A**, Overview of CHAD domain-containing proteins characterized in this study. The number of (no.) of CHAD proteins in the different kigdoms of life is indicated. **B**, Genetic loci of genes encoding CHAD domain-containing proteins (in black). Genes coding for polyP-metabolizing enzymes are highlighted in red. The DNA sequence upstream of RcCHAD is missing from contig RCOM_0386220 in the NCBI database. **C**, RT-PCR using primers binding to the *RcCHAD* coding sequence (left) results in a specific product amplified from cDNA from *Ricinus communis* leaves (right; DNA sequence in *SI Appendix* Fig. S2). **D**, Confocal microscopy of transgenic Arabidopsis lines expressing Ubi10p:RcCHAD-mCherry reveals RcCHAD to localize in the nucleus and nucleolus of root and hypocotyl cells (left, scale bars correspond to 20 μm). A western blot using an anti-mCherry antibody reveals a specific band at the predicted size of the RcCHAD-mCherry fusion protein (63 kDa). The Ponceau stained membrane is shown as loading control below (the major 55kDa band corresponds to RuBisCo).

To confirm if indeed RcCHAD is expressed in *Ricinus*, we performed RT-PCR experiments using *Ricinus* cDNA prepared from leave extracts. We detected a transcript corresponding to the predicted *RcCHAD* sequence (Fig. 1*C*) (*SI Appendix* Fig. S2). We next expressed RcCHAD carrying a C-terminal mCherry tag under the control of a constitutive promoter in the model plant *Arabidopsis thaliana*. We found that the fusion protein specifically localized to the nucleus and nucleolus in hypocotyl and root cells (Fig. 1*D*).

We next sought to determine crystal structures for different CHAD domains. Diffraction quality crystals developed for RcCHAD and CtCHAD, while crystals of SsCHAD diffracted only to ∼ 7 Å. Initial attempts to determine the RcCHAD structure using the molecular replacement method and the isolated CHAD domain of ygiF^221-422^ or an unpublished CtCHAD structure (PDB-ID 3e0s; both sharing ∼30% sequence identity with RcCHAD) failed (see Methods). We thus used the moderate anomalous signal present in the native RcCHAD dataset to locate a single Zn^2+^ ion. The structure was solved using the single-wavelength anomalous diffraction (SAD) method, and the refined model revealed a Zn^2+^ ion tetrahedrally coordinated by His50 and His136 originating from two symmetry-related RcCHAD molecules (*SI Appendix* Fig. S3). It has been previously speculated that CHAD domains may bind divalent cations using conserved histidine residues (39). We found however that His50 and His136 from the RcCHAD Zn^2+^ binding site are not conserved among other CHAD domains (*SI Appendix* Fig. S1), and consistently no metal ion binding sites were found in our crystal structures of CtCHAD (root mean square deviation, r.m.s.d. to the deposited PDB-ID 3e0s is ∼0.6 Å comparing 290 corresponding C_α_ atoms, *SI Appendix* Fig. S4) or ygiF (30).

We next compared the refined structures of the plant CHAD domain to the bacterial CtCHAD and ygiF structures. All CHAD domains fold into two four-helix bundles with up-down-up-down topology (Fig. 2*A*, *SI Appendix* Fig. S4). In all structures, the helical bundles are related by an almost perfect two-fold axis, and can be superimposed with r.m.s.d.’s ranging from 2.2-2.7 Å (*SI Appendix* Fig. S4). Notably, helices α4/α5 and α10/α11 are protruding the bundle cores in the RcCHAD and CtCHAD structures, forming a small pore in the center of the domain which we find to be absent in our ygiF structure (Fig. 2*B*, *SI Appendix* Fig. S4). This rationalizes the presence of ‘long’ (∼300 amino-acids, e.g. RcCHAD, CtCHAD) and ‘short’ (∼200 amino-acids, e.g. ygiF, SsCHAD) CHAD domains. Analysis of the surface charge distribution in the different CHAD domains revealed a highly basic central cavity, which is surrounded by two basic surface patches on each side (Fig. 2C). The basic amino-acids contributing to these surface patches are highly conserved among different CHAD proteins (*SI Appendix* Fig. S1). Similar, highly basic surface patches are present in many polyP metabolizing enzymes (*SI Appendix* Fig. S5). Analytical size-exclusion chromatography experiments indicated that the different CHAD proteins adopt different oligomeric states in solution, with ygiF and SsCHAD behaving as monomers, while RcCHAD and CtCHAD likely form dimers in solution (Fig. 2*D*, see *Discussion*).

**Figure 2.**
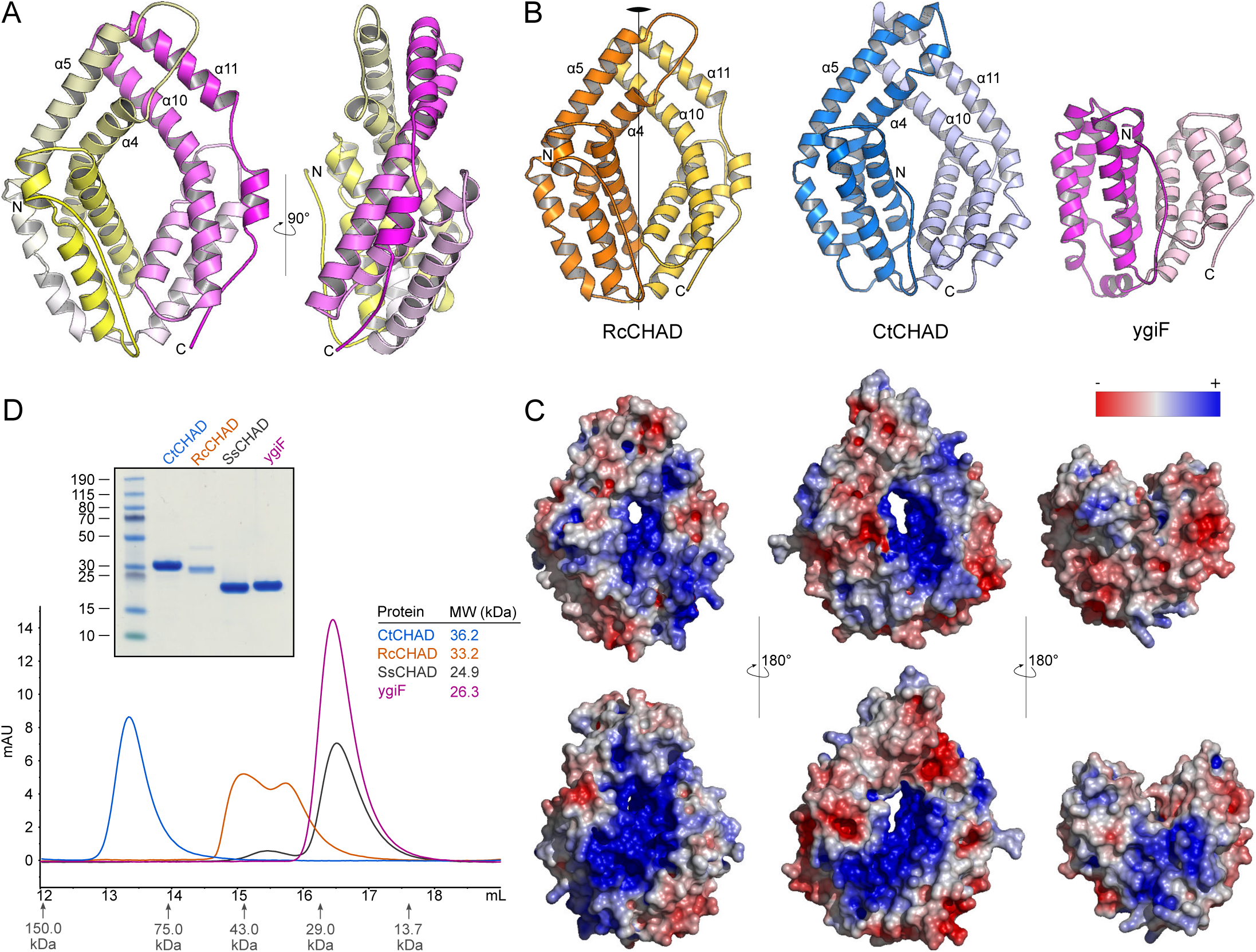
CHAD domains are helical bundles with two-fold internal symmetry and a conserved basic surface area. **A**, Overview of the CHAD domain architecture. Shown is a ribbon diagram of RcCHAD colored from N-(yellow) to C-terminus (magenta). **B**, Structural comparison of the RcCHAD, CtCHAD and ygiF CHAD domain (30) structures reveal the presence of two four-helix bundles in all CHAD domains, related by a pseudo two-fold symmetry (indicated by a vertical line). Note that the helices α4, α6, α10, and α11 in ygiF are much shorter when compared to RcCHAD and CtCHAD, and hence the central cavity is found open in ygiF. **C**, Identification of a conserved basic surface area in pro- and eukaryotic CHAD domains. Electrostatic potentials calculated in APBS (61) were mapped onto CHAD domain molecular surfaces in Pymol. Shown are front (upper panel) and back (lower panel) views. A highly basic surface area covers the front- and backside of the CHAD domain, and includes the central cavity present in RcCHAD and CtCHAD. **D**, Analytical size-exclusion chromatography reveals different oligomeric states for the CHAD domains analyzed in this study. An SDS-PAGE analysis of the respective peak fractions (pooled) is shown alongside.

Given their highly basic surface charge distribution, the fact that CHAD domains are found in polyP metabolizing enzymes or gene clusters (39), and that they can localize to polyP bodies (40), we next tested if CHAD domains directly bind polyPs.

We assayed polyP binding of RcCHAD, SsCHAD, CtCHAD and ygiF in quantitative grating-coupled interferometry (GCI) experiments (see Methods). Here, biotinylated polyP (chain length ∼100 P_i_ units) was coupled to the GCI chip, and the different proteins were used as analytes (*SI Appendix* Fig. S6). For our different CHAD domains, dissociation constants (K_*D*_) for polyP cover the nanomolar to the mid-micromolar range (Fig. 3*A*). The yeast polyP polymerase Vtc4p was used as a positive control (9), and BSA as negative control (Fig. 3*A*). RcCHAD and SsCHAD bind polyP with a one-to-one kinetics and with a *K*_*D*_ of 6 μM and 45 nM, respectively (Fig. 3*A*). In the case of CtCHAD and ygiF, the sensograms could not be explained by a simple one-to-one binding model. Instead, we observed two distinct association and dissociation events. A heterogeneous analyte model was used to fit the data (CtCHAD *K*_*D*_^1^=1.9 μM, *K*_*D*_^2^=147 nM; ygiF *K*_*D*_^1^=2.2 μM, *K*_*D*_^2^=40.5 μM). We performed competition experiments adding polyP (average chain length ∼7 P_i_ units) in various concentrations to a fixed concentration of CHAD protein sample used as analyte (Fig. 3*B*). In line with our direct binding assay, we find that polyP can efficiently compete for binding of SsCHAD and RcCHAD to the polyP-labeled surface of the GCI chip, with estimated IC_50_’s of ∼1 μM (Fig. 3*B*). In contrast, the highly negatively charged diadenosine pentaphosphate (AP5A) did not efficiently compete for binding of RcCHAD to polyP in GCI assays (Fig. 3*C*), and we could not observe detectable binding of SsCHAD or RcCHAD to GCI chips coated with biotinylated ssDNA (Fig. 3*D*). Together, our quantitative binding assays suggest that CHAD domains interact with polyPs with high specificity and selectivity. In line with this, we found that the presence of a C-terminal CHAD domain stimulated the previously reported tripolyphosphatase activity of ygiF (Fig. 3*E*).

**Figure 3.**
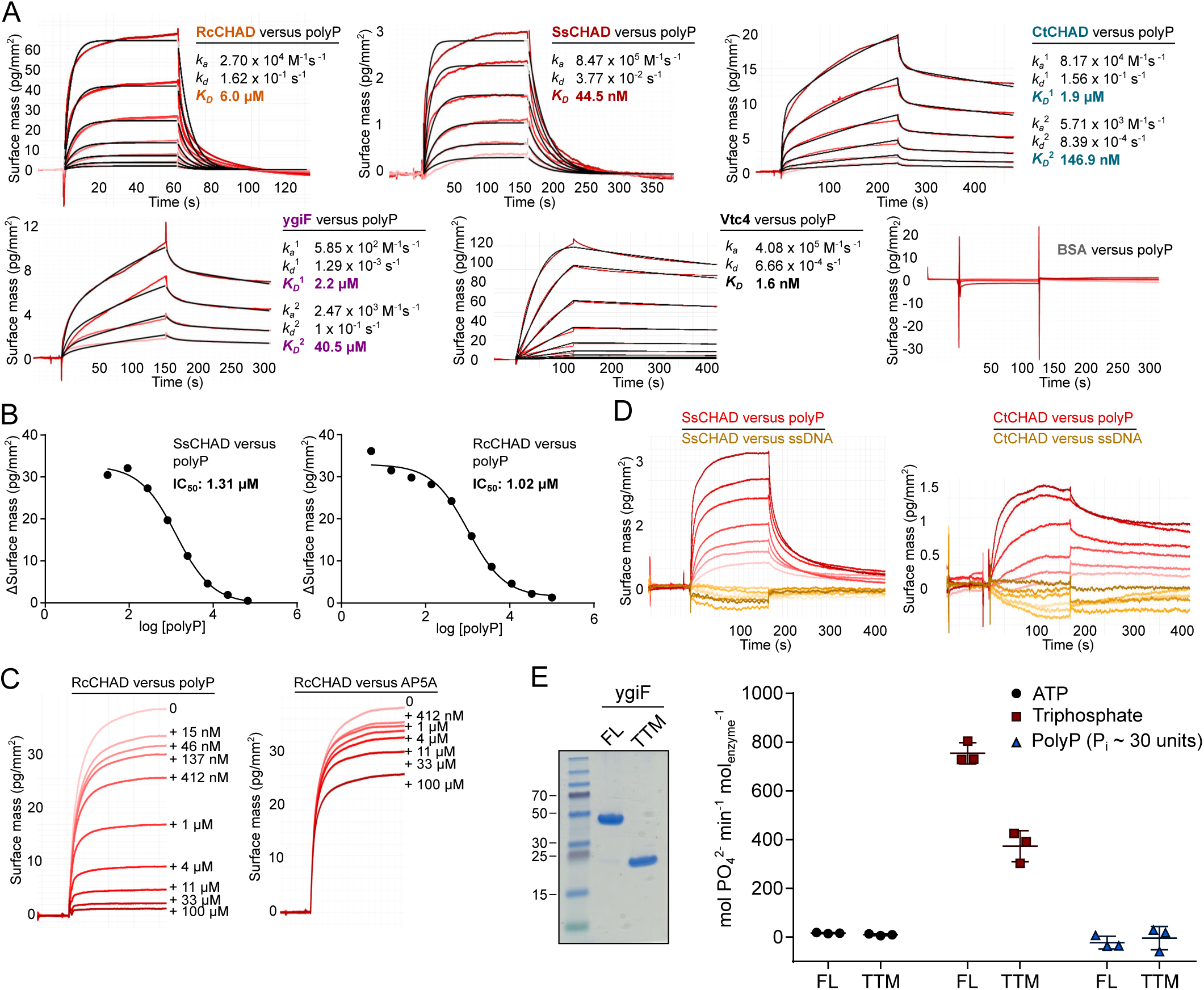
CHAD domains are specific polyP binding modules. **A**, Quantitative grating coupled interferometry (GCI) polyP binding assay. Biotinylated polyP (average chain-length is ∼ 100 Pi units) was immobilized on a streptavidin GCI chip, and the different CHAD domains were used as analytes (*SI Appendix* Fig. S6). The yeast polyP polymerase Vtc4 and BSA were used as positive and negative controls, respectively. Shown are recorded sensograms (in red) with the respective fits (in black), and including table summaries of the derived association rate constant (k_*a*_), dissociation rate constant (k_*d*_) and dissociation constant (K_*D*_). **B**, Short-chain polyPs (average chain-length ∼ 7 P_i_ units) can compete with CHAD domains for binding to the polyP-coated GCI chip. Shown are dose response curves with derived IC_50_ estimates. **C**, Diadenosine pentaphosphate (AP5A) cannot compete with RcCHAD for binding to the polyP-coated GCI chip (right panel) as efficiently as polyP (left panel, average chain-length ∼ 7 P _i_ units). Shown are sensograms of the association phase at indicated inhibitor concentration. **D**, Sensograms for SsCHAD and RcCHAD binding to biotinylated ssDNA (54 nt, in orange) or biotinylated polyP (average chain-length ∼ 100 P_i_ units, in red), respectively. **E**, Phosphohydrolase activities of ygiF-full length^1-433^ (FL) and ygif-TTM^1-200^ (TTM) vs. different phosphorylated substrates. Symbols represent raw data, lines indicate mean values, error bars denote the standard deviation of 3 independent replicates. An SDS-PAGE analysis of the purified proteins is shown alongside. The theoretical molecular weight is ∼22.3 kDa for TTM and ∼48.4 kDa for FL.

Next, we sought to identify residues in the CHAD domain involved in polyP coordination. A 1.7 Å structure of CtCHAD derived from crystals grown in 2 M (NH_4_)_2_SO_4_ revealed five sulfate ions bound in the central basic cavity of the CHAD domain (Fig. 4*A*). Crystals of CtCHAD grown in the presence of polyP (average chain length ∼7 P_i_ units) diffracted to 2.1 Å resolution and revealed continuous electron density transversing the central pore. The refined model includes a polyP 9-mer bound in the center of the CHAD domain and a tripolyphosphate moiety located at the distal side of the second basic surface patch (Fig. 4*B*). Additional differences electron density was too weak to be interpreted (dashed line in Fig. 4*B*). Superposition of the refined sulfate and polyP bound CtCHAD structures (r.m.s.d. is ∼ 0.5 Å comparing 304 corresponding C_α_ atoms) suggests that the sulfate ions mimic the positions of P_i_ units in the polyP chain binding across the CHAD domain center (Fig. 4*C*). The overall mode of polyP binding in CtCHAD is similar to the one seen in the previously reported Vtc4 – polyP and PPK2 – polyP complex structures (9, 24), with the polymer binding along a highly basic, solvent-exposed surface (Fig 4B, *SI Appendix* Fig. S5).

**Figure 4.**
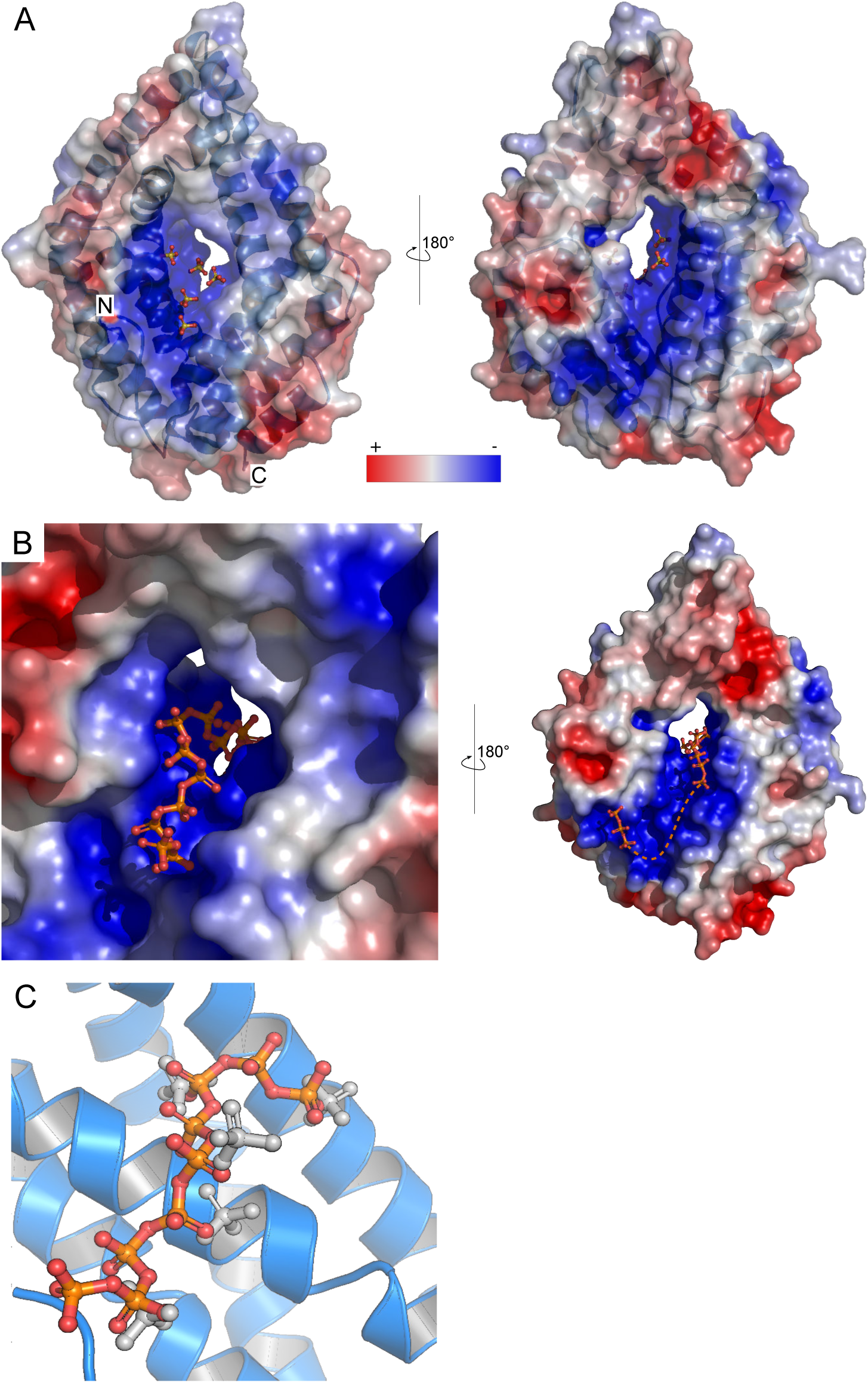
The basic surface area in CHAD domains provides a binding platform for polyPs. **A**, Sulfate ions (in bonds representation) originating from the crystallization buffer are bound to the basic surface area in CtCHAD. Shown are front and back views of CtCHAD as combined ribbon diagram and molecular surface. An electrostatic potential calculated in APBS has been mapped onto the molecular surfaces in Pymol. **B**, Overview of the polyP complex structure, obtained by crystallization of CtCHAD in the presence of 5 mM polyP (average length ∼7 P_i_ units). A polyP 9-mer and a tripolyphosphate moiety could be modeled (in bonds representation), with the polyP 9-mer occupying the central cavity and extending to both sides. The dashed line indicates the approximate position of several peaks in the F _o_-F_c_ difference electron density map, which however could not be modeled with confidence. **C**, Structural superposition of the sulfate ion- and polyP-bound CtCHAD structures (r.m.s.d. is ∼ 0.5 Å comparing 304 corresponding C_α_ atoms) reveals that the sulfate ions (in bonds representation, in gray) mimic the polyP 9-mer (in orange-red) in the CtCHAD-polyP complex.

We validated our structural model by mutational analysis of polyP interacting residues. In the CtCHAD – polyP complex structure, an apparent polyP 9-mer is coordinated by a set of conserved lysine and arginine residues lining the central cavity (Fig. 5*A*, *SI Appendix* Fig. S1). We mutated His253, Arg256 and Arg260, which form a hydrogen bonding network with polyP (Fig. 5*A*), to alanine. The mutant proteins bind polyP with 6-8 fold reduced affinity (Fig. 5*B*). Mutation of the corresponding residues His29, Arg32 and Arg36 in SsCHAD to alanine resulted in a ∼25 fold reduction in binding (Figs. 3A, 5B, *SI Appendix* Fig. S7). Additional mutation of Arg296 and Arg418/Tyr419 to alanine in CtCHAD led to ∼80-150 fold reduction in binding when compared to wild-type CtCHAD, while a His29/Arg32/Arg36/Arg69 SsCHAD quadruple mutant shows no detectable polyP binding in our GCI assay (Figs. 3*A*, 5*A,B*, *SI Appendix* Fig. S7). Together, these experiments reveal that the conserved basic amino-acids surrounding the central cavity in different CHAD proteins are involved in the specific recognition of phosphate polymers.

**Figure 5.**
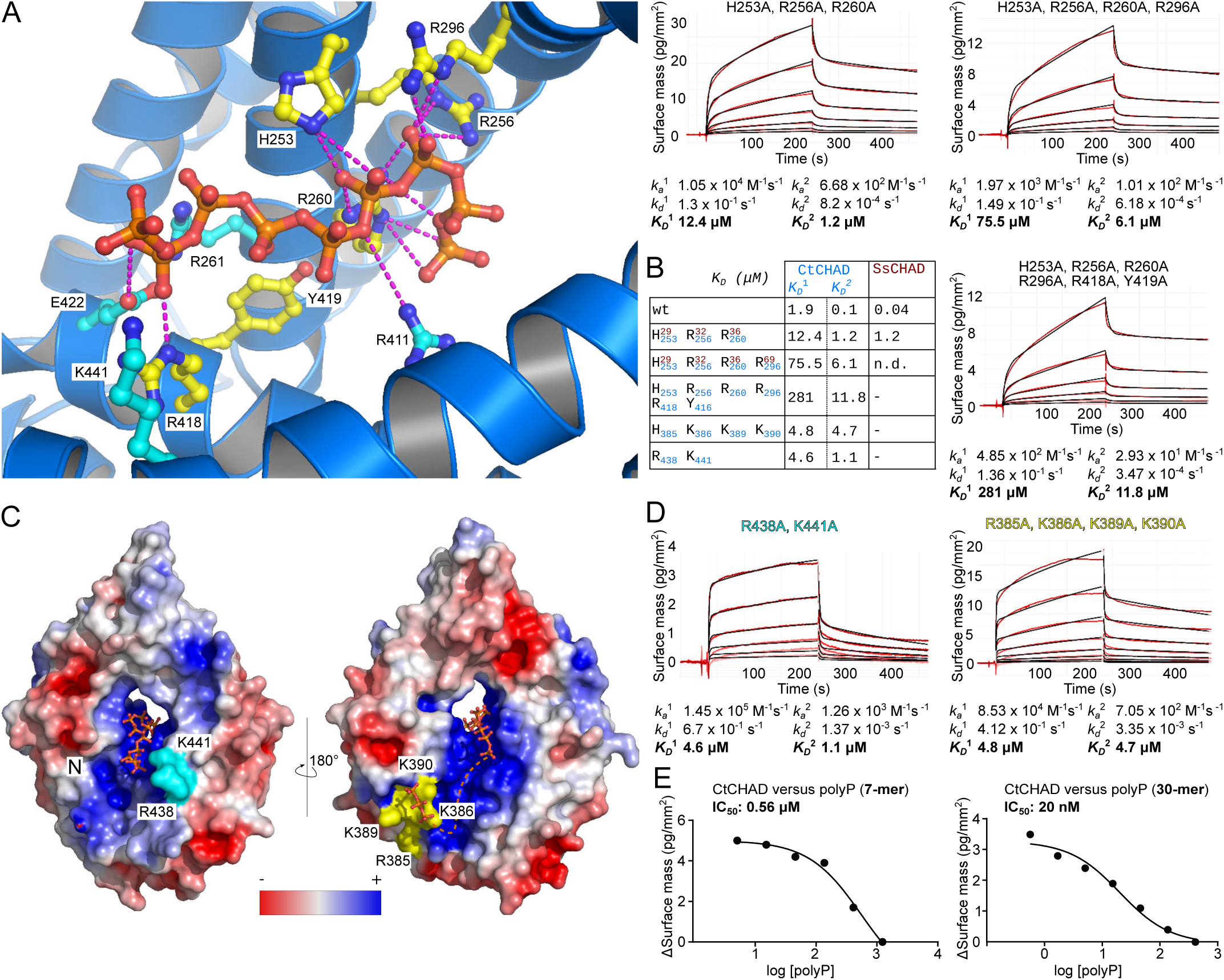
CtCHAD binds polyP through basic amino-acid residues distributed along the central cavity and the back of the protein. **A**, Detailed view of CtCHAD (blue ribbon diagram) bound to a polyP 9-mer (in orange, in bonds representation) and including selected conserved basic amino-acids involved in polyP binding (in cyan, residues included in mutational analyses shown in yellow). **B**, Mutations in the central basic binding surface in CtCHAD (dissociation constants in blue; corresponding mutations in SsCHAD in red) strongly decrease polyP binding in quantitative GCI assays. Shown are sensograms (in red), the respective fits (in black), and table summaries of the derived association rate constant (k _*a*_), dissociation rate constant (k_*d*_) and dissociation constant (K_*D*_). **C**, Identification of two distal polyP binding surfaces on the front side (shown in cyan) and back side (in yellow) of the CtCHAD – polyP complex structure. Mutations of con Alternatively, it is possible that one CHAD domain may simultaneously bind to several polyP chains immobilized on the GCI chip. In line with this, size-exclusion chromatography coupled to right-angle light scattering (SEC-RALS) revealed the presence of CtCHAD dimers and tetramers in the absence of polyP (*SI Appendix* Fig. S8). Addition of a long-chain polyP shifted CtCHAD into tetrameric and higher oligomeric states (*SI Appendix* Fig. S8). It is of not that CtCHAD also shows a crystal packing consistent with a dimer or tetramer, which would enable cooperative binding of several CHAD domains to a single polyP chain. In any case, our competition assays and our structure-based polyP binding site mutations in CtCHAD affected both binding kinetics,served arginine and lysine residues in both patches reduced polyP binding in GCI assays (**D**). **E**, GCI competition assays using short chain (average ∼ 7 P_i_ units) and long chain (average ∼ 30 P_i_ units) polyPs. Longer polyPs compete more efficiently (IC_50_ 20 nM versus 1 μM) with CtCHAD binding to the polyP coated GCI chip. Shown are dose-response curves with derived IC_50_ estimates.

Given that our analysis of different CHAD domain structures revealed the presence of additional, large and conserved basic surface patches, we next asked if these surfaces may be involved in the binding of long polyP chains. To this end, we generated additional point mutations in CtCHAD targeting either the ‘front’ or the ‘back’ side of the domain (shown in cyan and yellow in Fig. 5*C*, respectively). Mutation of the conserved Arg438 and Lys441 to alanine resulted in a ∼2-10 fold reduced binding affinity (Fig. 5*C,D*, *SI Appendix* Fig. S1). Mutation of Arg385, Lys386, Lys389 and Lys390 on the ‘back’ side of the domain had a similar effect (Fig. 5*C,D*). In line with this, longer polyPs (∼30 P_i_ units) compete much more efficiently (IC_50_ ∼ 20 nM) with CtCHAD binding to the polyP-coated (∼100 P_i_ units) GCI chip, when compared to polyP 7-mers (IC_50_ ∼ 1 μM) (Fig. 5*E*, *SI Appendix* Fig. S6). Together, these experiments suggest that CHAD domains can bind long polyP chains using their entire basic surface patch covering the ‘front’ and ‘back’ sides of the domain, as well as the central cavity/pore.

Our finding that CHAD domains can specifically bind polyPs with high affinity prompted us to further dissect the nuclear/nucleolar localization of RcCHAD stably expressed in Arabidopsis (Fig. 1D). It is presently unknown if polyPs are present in plants and where they would be localized. We transiently expressed RcCHAD-mCherry in tobacco leaves and again found the fusion protein to localize to the nucleus and to be further enriched in the nucleolus (Fig. 6). Next, we co-expressed RcCHAD-mCherry together with the bacterial polyP kinase EcPPK1. Expression of EcPPK1 has been previously reported to lead to cytosolic polyP accumulation in yeast cells (10). Notably, RcCHAD re-localized to punctate structures in the cytosol, which we assume to represent EcPPK1-generated polyP bodies (Fig. 6). Consistently, co-expression of RcCHAD-mCherry with a catalytically inactive variant of EcPPK1-mCitrine did not affect the nuclear/nucleolar localization of the CHAD domain in tobacco (Fig. 6). Based on these experiments, we speculate that RcCHAD may bind to a nucleolar/nuclear polyP pool in tobacco and in Arabidopsis, and that PPK1-generated polyPs may force RcCHAD to relocate to the cytosol.

**Figure 6.**
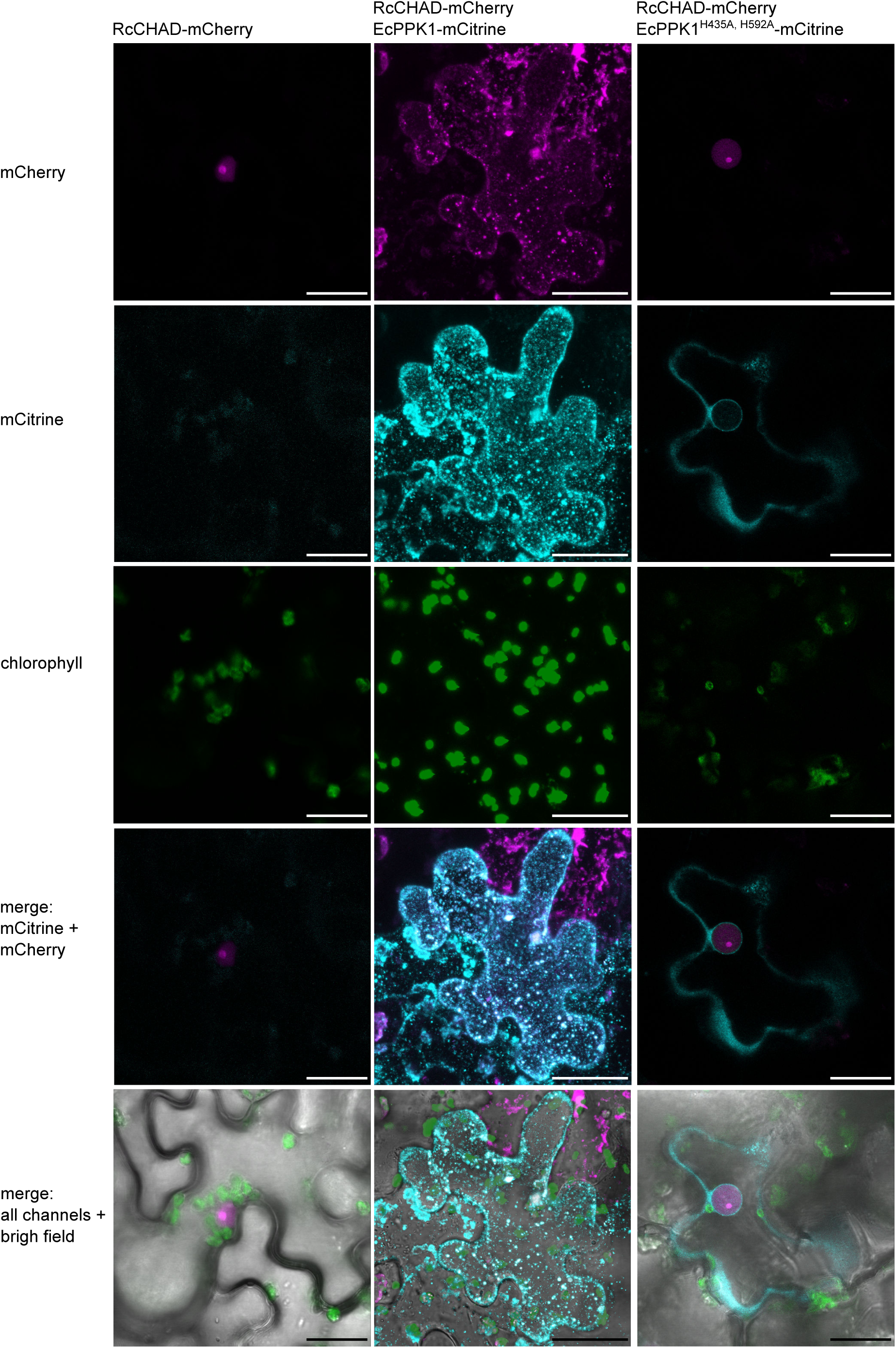
RcCHAD localizes to the nucleus and nucleolus of tobacco cells, and co-localizes with EcPPK1 to EcPPK1-generated polyP granules. Transient expression of Ubi10p:RcCHAD-mCherry in tobacco leaves reveals a nuclear/nucleolar localization of the fusion protein (left). Expression of Ubi10p:EcPPK1-mCitrine induces the formation of polyP granules (center), not observed when using a catalytically impaired version of the enzyme (EcPPK1^H435A, H592A^, right). RcCHAD co-localizes with EcPPK1-generated granules (center). Shown are images obtained through confocal microscopy of tobacco leaves transiently expressing the indicated constructs, scale bars correspond to 50 μm.

## Discussion

CHAD domains have been originally defined as ‘conserved histidine α-helical domains’, with the histidines acting as metal chelators and/or phosphoacceptors (39). While we found a Zn^2+^ ion coordinated by two histidine residues in our RcCHAD structure (*SI Appendix* Fig. S3), the contributing histidines are not conserved among CHAD family proteins (*SI Appendix* Fig. S1), and no metal ions could be located in our CtCHAD and ygiF structures (30). This makes it unlikely that CHAD domains are metal binding proteins. Our structural analysis revealed that CHAD domains are helical bundles featuring an unusual internal symmetry. A set of highly conserved basic amino-acids contributes to the formation of a large basic surface area, which has evolved to sense long-chain polyPs, but not DNA or other P_i_ containing ligands. Our plant, archaeal and bacterial CHAD proteins bind polyPs with dissociation constants in the μM to nM range, in good agreement with the cellular concentrations reported for polyPs in different organisms (28, 41, 42). Our quantitative biochemical experiments show that GCI can be employed to quantify polyP binding to CHAD domains. In contrast to other methods, in GCI assays the heterogeneous chain length of the polyP ligand does not affect the accuracy of the derived kinetic parameters. The sensograms of CtCHAD and ygiF binding to polyP could only be explained using a heterogeneous analyte model (Fig. 3*A*). We speculate that different oligomeric states observed with the CHAD domains used in this study may account for this behavior (Fig. 2*D*). Alternatively, it is possible that one CHAD domain may simultaneously bind to several polyP chains immobilized on the GCI chip. In line with this, size-exclusion chromatography coupled to right-angle light scattering (SEC-RALS) revealed the presence of CtCHAD dimers and tetramers in the absence of polyP (*SI Appendix* Fig. S8). Addition of a long-chain polyP shifted CtCHAD into tetrameric and higher oligomeric states (*SI Appendix* Fig. S8). It is of note that CtCHAD also shows a crystal packing consistent with a dimer or tetramer, which would enable cooperative binding of several CHAD domains to a single polyP chain. In any case, our competition assays and our structure-based polyP binding site mutations in CtCHAD affected both binding kinetics, suggesting that our reported dissociation constants represent *bona fide* polyP binding events (Fig. 5*B*). Taken together, our binding assays and our polyP complex structure suggest that CHAD domains are polyP-binding modules that lack enzymatic activity (30).

The conserved structural, biochemical and sequence features of bacterial, archaeal and eukaryotic CHAD domains suggest that these polyP-binding modules may have a common, ancient evolutionary origin. We identified an expressed CHAD domain-containing protein in the plant *Ricinus comunis* L. (RcCHAD) which represents the only CHAD domain currently known in plants. Interestingly, RcCHAD is ∼ 80 % sequence identical to a protein from the rhizosphere bacterium *Duganella* (*43*) (*SI Appendix* Fig. S1). Thus, *RcCHAD* might have been acquired by the plant via horizontal gene transfer from a soil-living bacterium.

While we have biochemically characterized CHAD domains as polyP-binding proteins, their physiological roles remain to be defined. It has been previously shown that CHAD domain-containing proteins localize to polyP bodies in the bacterium *Ralstonia eutropha*, and that their over-expression can relocalize polyP granules to the cell poles (40). However, genetic depletion of Ralstonia CHAD proteins or of ygiF did not result in any apparent phenotype (38, 40). Our enzymatic assays indicate that CHAD domains may assist polyP-metabolizing enzymes in recruiting their substrates. We speculate that the large polyP binding surface in CHAD and its central cavity/pore (which is occupied by a polyP polymer in our complex structure, Fig. 4*B*) may render fused polyP-metabolizing enzymes highly processive, as previously speculated (44). Consistently, about half of the annotated CHAD-containing proteins harbor N-terminal CYTH/TTM domains, which we and others have previously characterized as short-chain inorganic polyphosphatases (30, 36, 38).

In bacteria, polyP granules are spatially restricted in the nucleoid region (3). PolyP has also been shown to accumulate in the nucleolus of myeloma cells (45). We could observe a specific nuclear/nucleolar localization of RcCHAD in different cells and tissues when stably expressed in Arabidopsis, or transiently expressed in tobacco. We infer from this finding that polyP in plants may be located in the nucleolar compartment, as reported for animal cells (45). In line with this, ectopic expression of PPK1 leads to a relocalisation of RcCHAD to presumed polyP granules in the cytosol. The polyP binding domain from *E. coli* PPX1 has been previously used to detect polyP pools in fungal, algal and mammalian cells (46, 37, 47, 45). Based on its small size, high polyP binding affinity and specificity together with the well characterized polyP binding mechanism, we now propose CHAD domains as molecular probes to dissect polyP metabolism and storage in pro- and eukaryotic cells.

## Acknowledgements

We thank staff at beam-line X06DA – PXIII of the Swiss Light Source (SLS), Villigen, Switzerland for technical help during data collection, and F. Spiga for advice on the GCI experiments. This work was supported by an ERC starting grant from the European Research Council under the European Union’s Seventh Framework Programme (FP/2007-2013) / ERC Grant Agreement n. 310856 and the Howard Hughes Medical Institute (International Research Scholar Award; both to M.H.)

## Author Contributions

L. L-O. and M. H. designed the study, L. L-O. performed the majority of the experiments and analyzed data. U.H. and L.L-O. developed the GCI binding assay. L.L-O. and J.Z. performed cell biology experiments. M.H. and L.L-O. analyzed he crystallographic data. L. L-O. and M. H. wrote the manuscript with input from the other authors.

## Methods

### Reverse transcription polymerase chain reaction (RT-PCR)

RNA was extracted from ∼ 100 mg of *Ricinus communis* leaves with the Rneasy Plant Mini Kit (Qiagen). 2 μg of RNA was treated with Dnase I (Qiagen), copied to cDNA using an Oligo dT and the SuperScript™ II Reverse Transcriptase (Invitrogen). RT-PCR was performed with primers RcCHAD_RT_F (5-ATTGCCCAGGCAAAGCGTCATGC-3) and RcCHAD_RT_R (5-TTAGTGACGTAACTGTGGTGC-3). The RT-PCR product was resolved on a 0.8 % agarose gel, revealing a single DNA product. The band was excised and sequenced using the RcCHAD_RT_F/R primers. Sequencing results were analyzed using CLC Main Workbench 7.9.1 (Qiagen).

### Generation of Arabidopsis transgenic lines

The *RcCHAD* coding sequence was cloned in pDONR221, the *UBIQUITIN10* promoter (pUBI10) in pDONR P4-P1R, and the mCherry fluorescence tag in pDONR P2R-P3, using the Gateway™ BP Clonase™ II Enzyme mix (Merck). Constructs were assembled with the Gateway™ LR Clonase™ enzyme mix (Merck) into the vector pH7m34GW (48). *Agrobacterium tumefaciens* (pGV2260) was transformed with pH7m34GW harboring the construct pUBI10:RcCHAD-mCherry. *Arabidopsis thaliana* was transformed using the floral dip method (49), and plants were selected in 1/2 MS medium (1/2 MS [Duchefa], 1 % [w/v] sucrose, 0.5 g/L MES pH 5.7, 0.8 % agar), supplemented with 20 μg/mL hygromycin.

### Transient protein expression in *Nicotiana benthamiana* leaves

The EcPPK1 coding sequence (uniprot ID C3T032) was cloned in pDONR221. PPK1 enzyme-dead mutations (H435A/H592A) were introduced by site-directed mutagenesis. EcPPK1 constructs were assembled together with Ubi10p (in pDONR P4-P1R) and mCitrine (in pDONR P2R-P3) into the pH7m34GW vector using the Gateway™ BP Clonase™ II Enzyme mix (Merck). *Agrobacterium tumefaciens* was transformed with Ubi10p:RcCHAD-mCherry, Ubi10p:EcPPK1-mCitrine, Ubi10p: EcPPK1^[H435A,H592A]^-mCitrine and p19. For each construct, 10 mL of *Agrobacterium* culture, grown overnight at 28 °C, was collected by centrifugation. Cells were resuspended in 10 mL of infiltration solution (10 mM MgCl_2_, 10 mM MES pH 5.6 and 100 mM acetosyringon) and incubated for 3 h in darkness. For co-localization experiments, cells expressing two different constructs were mixed in equal volumes before infiltration. Cells expressing p19 were added to all solutions. Tobacco leaves were infiltrated using a 0.5 mL syringe and plants were imaged after 2 d.

### Confocal microscopy

7 d old *Arabidopsis* T3 seedlings expressing pUBI10:RcCHAD-mCherry, or tobacco leaves infiltrated with pUBI10:RcCHAD-mCherry and/or pUBI10:EcPPK1-mCitrine were imaged using a LSM 780 confocal microscope (Zeiss) equipped with a 40x NA 1.2 water C-Apochromat lens. Transmission was imaged at 514nm. mCitrine and mCherry were imaged using a GaAsP detector upon excitation at 514nm and 594nm, respectively, and emission between 517-552 nm (mCitrine) and 606-632 nm (mCherry). Chlorophyll was imaged with a PMT detector upon excitation at 594nm and with emission between 653-658 nm. Images were analyzed using Fiji (50).

### Western blotting

Arabidopsis seedlings were snap-frozen in liquid nitrogen and homogenized with mortar and pestle. The plant extract was resuspended in 50 mM Tris pH 8.0, 150 mM NaCl, 0.5 % (v/v) Triton X-100 and cOmplete^™^ EDTA-free Protease Inhibitor Cocktail (Merck). 50 μg of protein extract (estimated by Bradford, Bio-Rad), pre-boiled for 5 min, was run on a 10 % SDS-PAGE gel. Blotting was done on a nitrocellulose membrane (GE Healthcare). After blocking with TBS buffer supplemented with 0.1 % (v/v) Tween 20 and 5 % (w/v) powder milk, the membrane was first incubated for 1 h with an anti-mCherry antibody (Abcam, ab167453, dilution 1:2000), and then with an anti-rabbit peroxidase conjugate antibody (Calbiochem, dilution 1:10,000, 1 h). RuBisCO proteins were visualized with Ponceau (0.1 % [w/v] Ponceau S in 5 % [v/v] acetic acid) as loading controls.

### Protein expression and purification

The coding sequences of RcCHAD (uniprot ID B9T8Q5), SsCHAD (uniprot ID Q97YW1), and CtCHAD^208-522^ (uniprot ID Q8KE09) were obtained as synthetic genes from Geneart (Life Technologies) and cloned into the vector pMH-HT (providing a N-terminal 6xHis tag followed by a tobacco etch virus (TEV) protease cleavage site) by Gibson (51) or restriction-based cloning. Plasmids were transformed in *Escherichia coli* BL21 (DE3) RIL cells. For protein expression, cells were grown in terrific broth (TB) medium at 37 °C until OD_600_ ∼ 0.6, induced with 0.25 mM isopropyl β-d-galactoside (IPTG), and grown at 16 °C for ∼ 16 h. Cell pellets were collected by centrifugation at 4,500 x g for 30 min, resuspended in lysis buffer (50 mM sodium phosphate pH 7.5, 500 mM NaCl, lysozyme, Dnase I and cOmplete™ Protease Inhibitor Cocktail [Merck]), and disrupted by sonication. The cell suspension was spun down at 18,000 x g for 1 h, and the supernatant was loaded onto a Ni^2+^ affinity column (HisTrap HP 5 ml, GE Healthcare). The column was washed with 5 column volumes (CV) of buffer A (50 mM PBS pH 7.5, 500 mM NaCl), 5 CV buffer B (50 mM PBS pH 7.5, 1 M NaCl), and 5 CV buffer C (250 mM PBS pH 7.5, 500 mM NaCl). Proteins were eluted with buffer A supplemented with 0.5 M imidazole pH 8.0, and cleaved overnight with TEV at 4 °C during dialysis in buffer A. RcCHAD was further purified by cation exchange (HiTrap SP HP cation exchange chromatography column, GE Healthcare), CtCHAD by size-exclusion chromatography, and SsCHAD by a second Ni^2+^ affinity step. All samples were purified to homogeneity by size-exclusion chromatography on a Superdex 75 HR26/60 (GE Healthcare) equilibrated in buffer A. Protein concentrations were estimated by ultraviolet absorption at 280 nm (A_280nm_) and using the respective theoretical molecular extinction coefficient calculated with the program PROTPARAM (https://web.expasy.org/protparam/). Mutations were introduced by site-directed mutagenesis, mutant proteins were purified like wild-type. Vtc4^189-487^, ygiF-full length^1-433^, ygiF-TTM^1-200^ and ygiF-CHAD^201-433^ were purified as described previously (9, 30).

### Crystallization and crystallographic data collection

Hexagonal RcCHAD crystals developed at room temperature from hanging drops containing 1.5 μL of protein solution (1.6 mg/mL) and 1.5 μL of crystallization buffer (3 M NaCl, 0.1 M Bis-Tris pH 6.0) suspended over 0.5 mL of crystallization buffer as reservoir solution. Orthorhombic CtCHAD crystals grew at room temperature in hanging drops containing 1.5 μL of protein (8 mg/mL) and 1.5 μL of crystallization buffer (2 M (NH_4_)_2_SO_4_, 5 % isopropanol). The CtCHAD – polyP complex was prepared by mixing CtCHAD at 8 mg/mL with short chain polyP (BK Giulini GmbH, Calgon® 188, average chain length ∼ 7 P_i_ units) to a final concentration of ∼ 5 mM. Crystals developed in 0.4 M (NH_4_)_3_PO_4_ in hanging drop setups. All crystals were cryo-protected by serial transfer into crystallization buffer supplemented with 20-30 % ethylene glycol, and snap-frozen in liquid nitrogen.

### Crystallographic data collection, structure solution and refinement

Diffraction data were collected at beam line X06DA – PXIII of the Swiss Light Source (SLS), Villigen, Switzerland and data processing and scaling was done in XDS (version Jan 26, 2018) (52). For RcCHAD, a complete dataset at 2.3 Å resolution containing a weak anomalous signal to ∼6 Å resolution (λ ∼ 1.0 Å) was recorded (*SI Appendix* Table. S1). A single Zn^2+^ ion was located by SHELXD (53) and the structure was solved using the single-wavelength anomalous dispersion (SAD) method as implemented in the program phenix.autosol (54). The resulting model was completed in alternating cycles of manual model correction in the program COOT (55) and restrained refinement in autoBUSTER (Global Phasing Ltd.). The sulfate ion-bound structure of CtCHAD was solved to 1.7 Å resolution using the molecular replacement method as implemented in PHASER (56) (PDB-ID 3E0S was used as search model), and refined in phenix.refine (54). Isomorphous crystals of the CtCHAD – polyP complex diffracted to 2.1 Å resolution, restraints for a polyP 9-mer were generated using the program JLigand (57) and the structure was refined in REFMAC5 (58). Structural representations were done in Pymol (http://pymol.sourceforge.net/) and using the ray tracer POVRAY (http://www.povray.org/). Secondary structure assignments were calculated with DSSP (59).

### Analytical size-exclusion chromatography

Gel filtration experiments were performed using a Superdex 200 Increase 10/300 GL column (GE Healthcare) pre-equilibrated in buffer A. 500 μL of the respective protein (0.5 mg/mL) was loaded sequentially onto the column and elution at 0.75 ml/mL was monitored by ultraviolet absorbance at 280 nm. Peak fractions were analyzed by SDS-PAGE gel electrophoresis.

### Grating-coupled interferometry (GCI) binding assays

GCI assays were performed using a Creoptix WAVE system (Creoptix AG) (60) as illustrated in *SI Appendix* Fig. S6. Experiments were performed using 4PCP WAVE GCI chips (quasi-planar polycarboxylate surface; Creoptix AG). After conditioning with borate buffer (100 mM sodium borate pH 9.0, 1 M NaCl), the chip was immobilized in all channels with streptavidine and BSA via a standard amine-coupling: activation with 1:1 mix of 400 mM N-(3-dimethylaminopropyl)-N’-ethylcarbodiimide hydrochloride and 100 mM N-hydroxysuccinimide, immobilization with 30 μg/mL of streptavidine in 10 mM sodium acetate pH 5.0, passivation with 5 % BSA in 10 mM sodium acetate pH 5.0, and quenching with 1 M ethanolamine pH 8.0. We did not succeed in coupling the CHAD domains directly to the chip using various methods. Hence, biotinylated medium chain polyP (5-20 ug/uL, Kerafast) or 5’-biotinylated single-strand DNA (5 ug/uL, 54 nucleotides, Metabion) was bound to the chip surface. Analytes were injected in a 1:2 dilution series in 50 mM Bis-Tris pH 7.5, 150 mM NaCl at 25 °C. Blank injections (every 3 cycles) were used for double referencing, and a DMSO calibration curve (0-2 % DMSO, 4 dilutions) for bulk correction. Data analysis was performed using the Creoptix WAVEcontrol software version 3.5.13 (applied corrections: X and Y offset, DMSO calibration, double referencing) and a one-to-one binding model was used to fit all experiments with the exception of CtCHAD and ygiF, in which we used a heterogeneous analyte model. For competition experiments, fixed concentration of CHAD proteins were incubated with a dilution series of short-chain polyP (average chain length ∼ 7 P_i_ units, BK Giulini GmbH, Calgon® 188), polyP (average chain length ∼ 30 P_i_ units, BK Giulini GmbH, Calgon® 322) or P^1^,P^5^-Di(adenosine-5′) pentaphosphate pentasodium salt (Merck).

### Analysis of oligomeric states by means of Right Angle Static Light Scattering (RALS)

The oligomeric states of ‘apo’ and polyp-loaded CtCHAD were analyzed by size exclusion chromatography (SEC) combined with right angle light scattering (RALS) using a OMNISEC RESOLVE/REVEAL combo, providing a GPC/SEC tetra-detector. Instrument constants were determined with defined concentrations of BSA. Apo CtCHAD, or CtCHAD incubated with 1 mM polyP (average chain length ∼ 30 P_i_ units, BK Giulini GmbH, Calgon® 322) at room temperature for 3 h, were analyzed in aliquots of 50 μl each at a sample concentration of 2 mg/ml (in 20 mM Hepes pH 7.5, 150 mM NaCl) on a Superdex 200 10/300 increase column (GE Healthcare, Germany) at a column temperature of 35 °C and a flow rate of 0.7 mL/min. Samples were analyzed using the OMNISEC software, version 10.41 (*SI Appendix* Fig. S8).

### Phosphohydrolase activity measurements

10 nM of ygiF-full length^1-433^ and ygiF-TTM^1-200^ were incubated for 7 min at 37 °C with 500 μM of substrate in reaction buffer (20 mm Bis-Tris propane pH 8.5, 150 mm NaCl, 5 mm MgCl_2_). The substrates tested were ATP (Merck), sodium tripolyphosphate (Merck), and polyP (average length ∼ 30 P_i_ units, BK Giulini GmbH, Calgon® 322). 100 μL of the reaction was incubated for 5 min with 28 μL of a malachite green solution containing 3 mM malachite green, 15 % (v/v) sulfuric acid, 1.5 % molybdate (w/v) and 0.2% (v/v) Tween 20. The absorption at A_595nm_ was measured using a synergy H4 plate reader (Biotek). Blanks were obtained for each substrate, by adding heat-inactivated enzyme (boiled for 5 min at 95 °C) to the respective reactions. Experiments were performed in triplicates.

## Figure legends

**SI Appendix Fig. S1.**
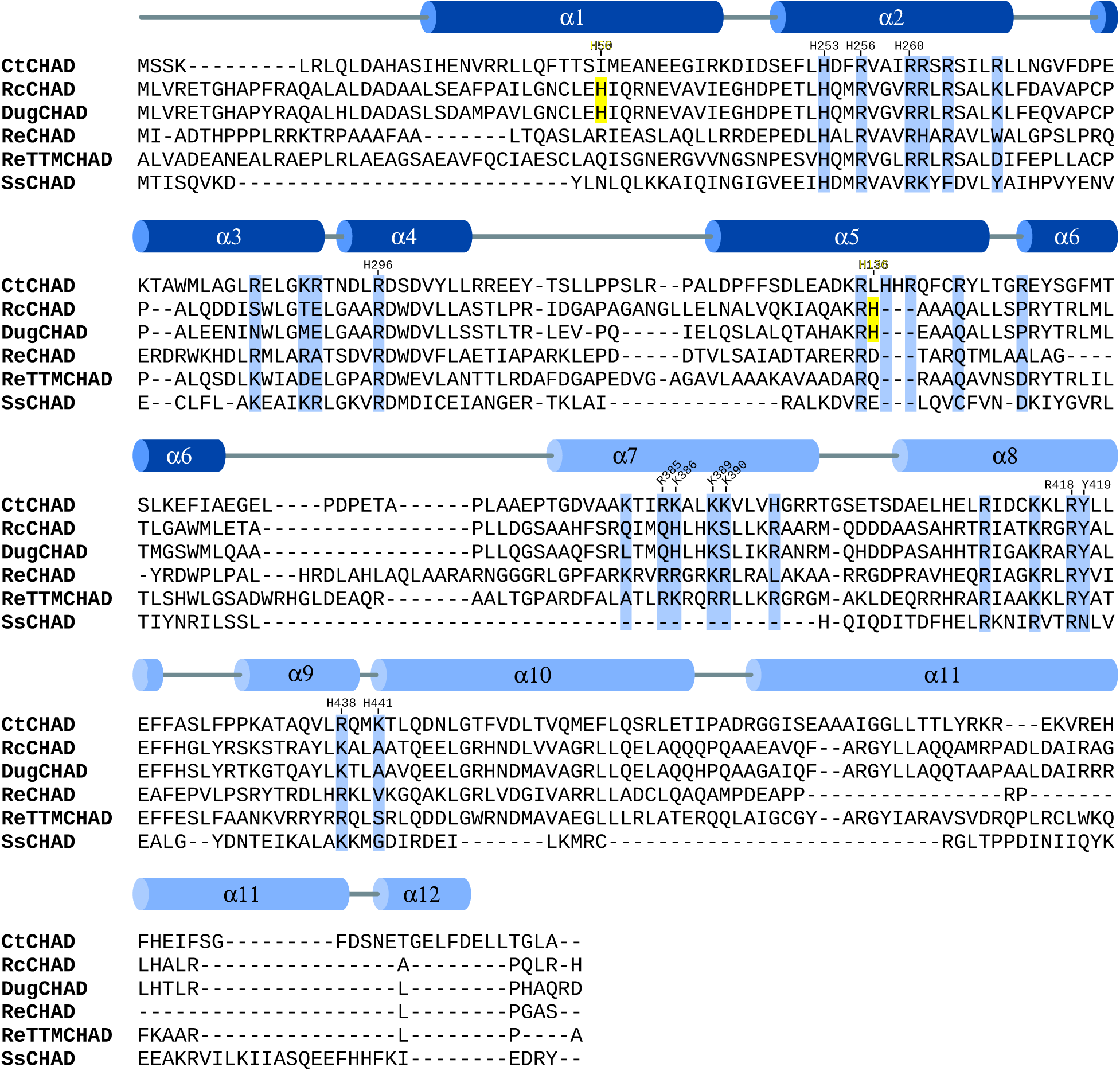
Sequence alignment of CHAD domains. Structure-based sequence alignment of CtCHAD (*Chlorobium tepidum*, unitprot ID Q8KE09), RcCHAD (*Ricinus communis*, B9T8Q5), DugCHAD (*Duganella* sp., A0A1H7Y2Q3), ReCHAD (*Ralstonia eutropha*, H16_A0104), ReTTMCHAD (*Ralstonia eutropha*, H16_B1017) and SsCHAD (*Sulfolobus solfataricus*, Q97YW1) and including a secondary structure assignment for CtCHAD calculated with DSSP (59). α-helices belonging to the two four-helix bundles are colored in dark and light blue, respectively. The histidines involved in Zn^2+^ ion coordination in the RcCHAD structure are highlighted in yellow, conserved basic residues contributing to the polyP binding surface are shown in blue, numbers indicate the positions of residues in CtCHAD used in the mutational analysis in Figure 5.

**SI Appendix Fig. S2.**
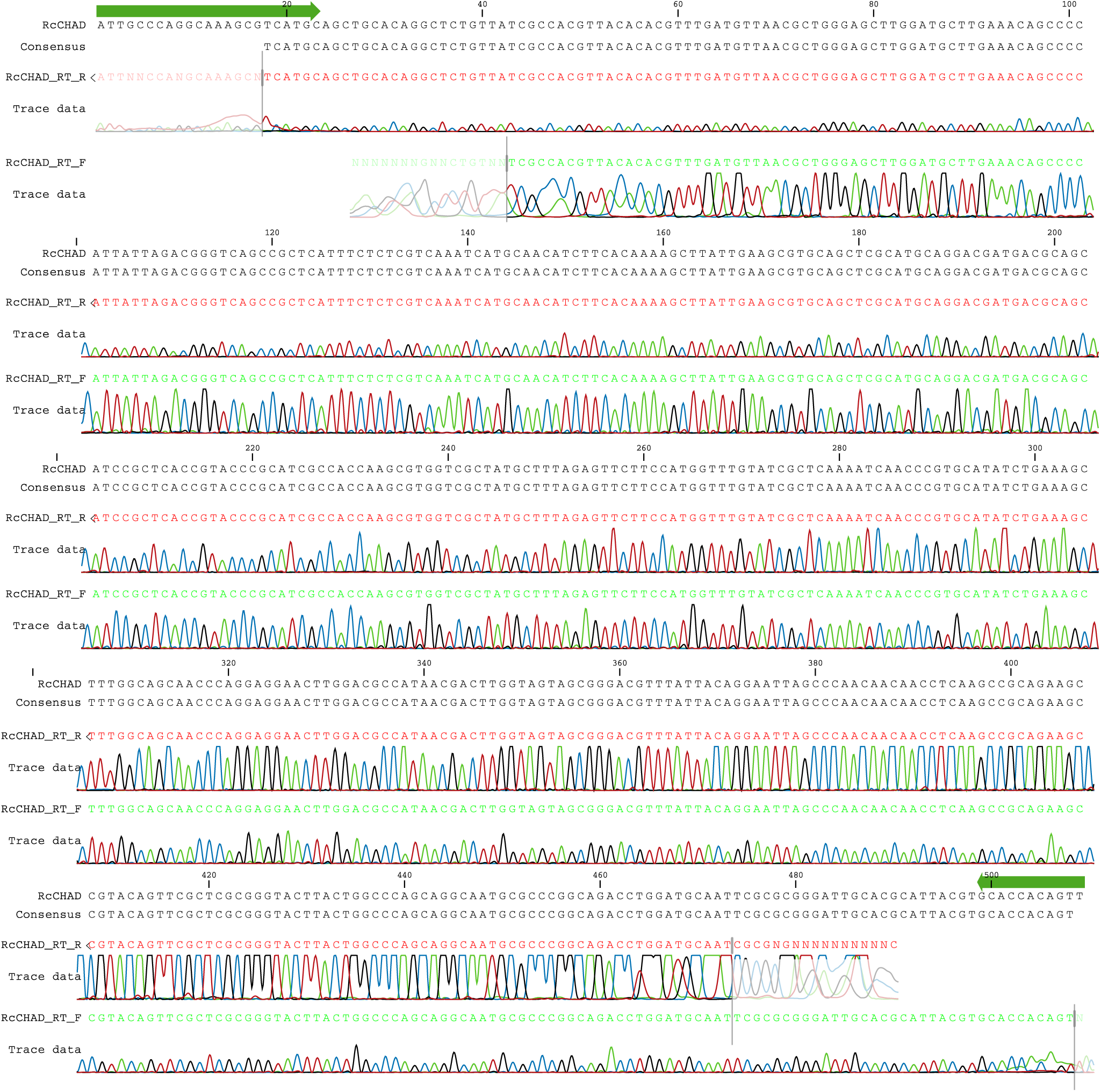
Sequencing results of the RT-PCR product amplified from *Ricinus* cDNA using *RcCHAD*-specific primers. Primers are shown in green, the obtained sequence matches the NCBI contig ID RCOM_0386220.

**SI Appendix Fig. S3.**
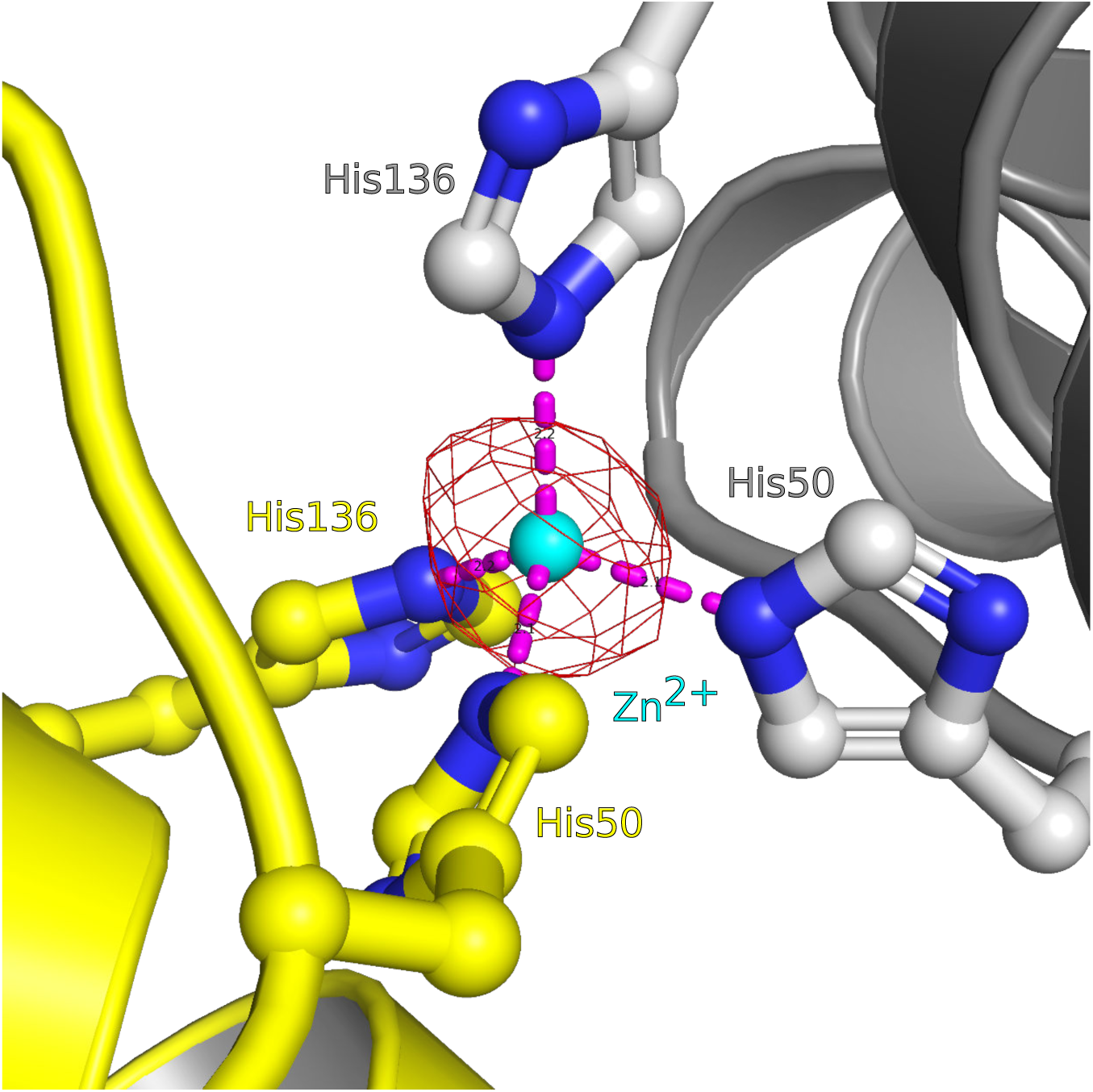
Close-up view of the Zn^2+^ binding site located in the RcCHAD structure. The binding site around the Zn^2+^ ion (cyan sphere) is formed by the non-conserved His50 and His136 (in bonds representation, interactions shown as dotted lines) at the interface of two symmetry-related molecules (shown in yellow and gray, respectively). A phased omit difference density map contoured at 8 s is shown alongside (red mesh).

**SI Appendix Fig. S4.**
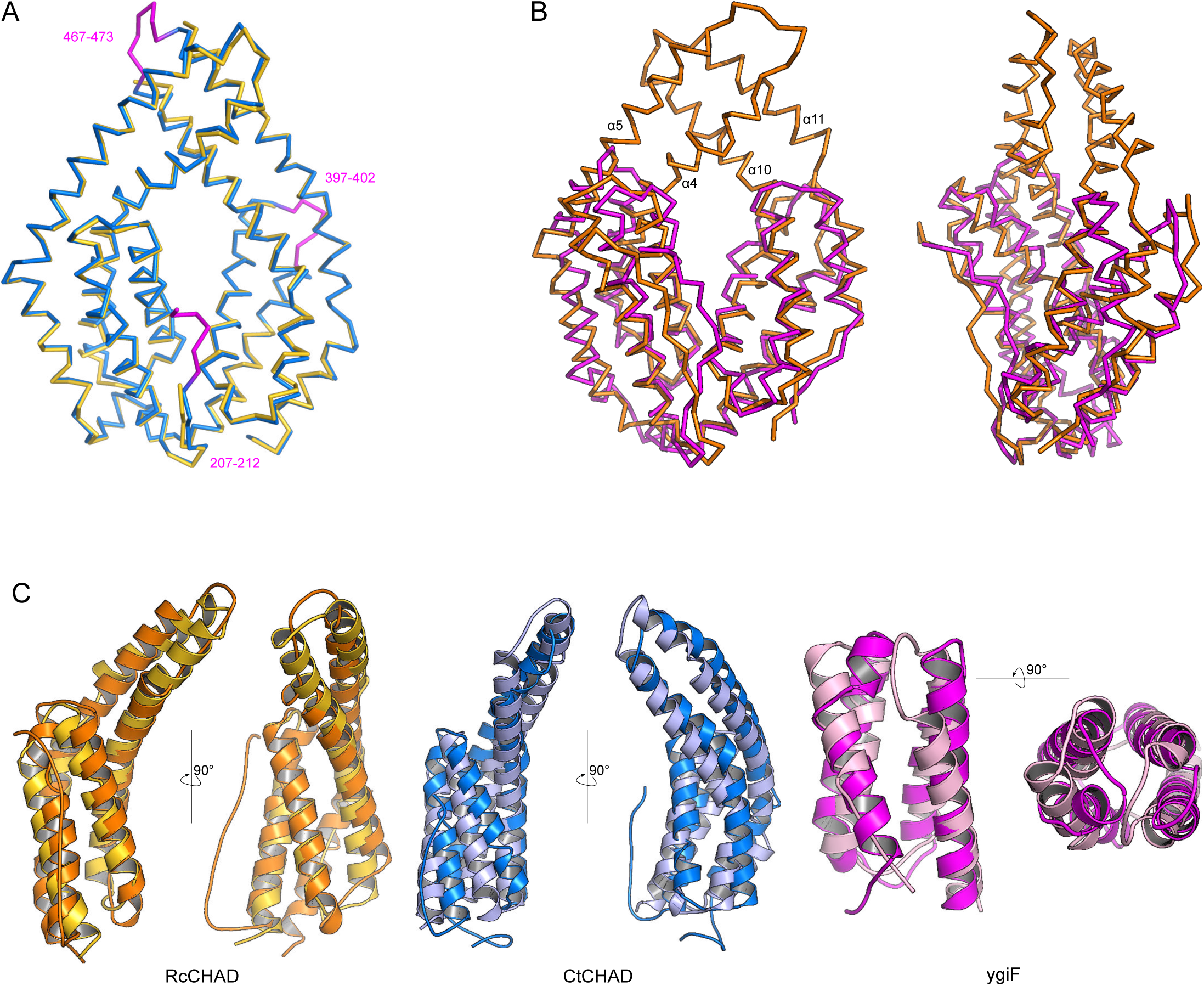
CHAD domains are formed by two four-helix bundles related by a pseudo two-fold internal symmetry. **A**, Structural superposition of our CtCHAD structure (in blue) with a second CtCHAD crystal form (PDB-ID 3e0s, in yellow; r.m.s.d. is ∼ 0.6 Å comparing 290 corresponding C_α_ atoms). Shown are C_α_ traces, with loop regions present in our structure and missing in PDB-ID 3e0s highlighted in magenta. **B**, Structural superposition of RcCHAD (in orange) with the CHAD domain from E. coli ygiF (PDB-ID 5a61, residues 205-428, in magenta) reveals the presence of protruding helices in RcCHAD closing the central cavity (r.m.s.d. is ∼ 2.4 Å comparing 184 corresponding C_α_ atoms). **C**, Structural superposition of the two four-helix bundles in RcCHAD (residues: 7-166 versus 167-303), CtCHAD (residues: 207-373 versus 374-522) and ygiF (residues: 205-336 versus 337-428) shown as ribbon diagrams in two orientations.

**SI Appendix Fig. S5.**
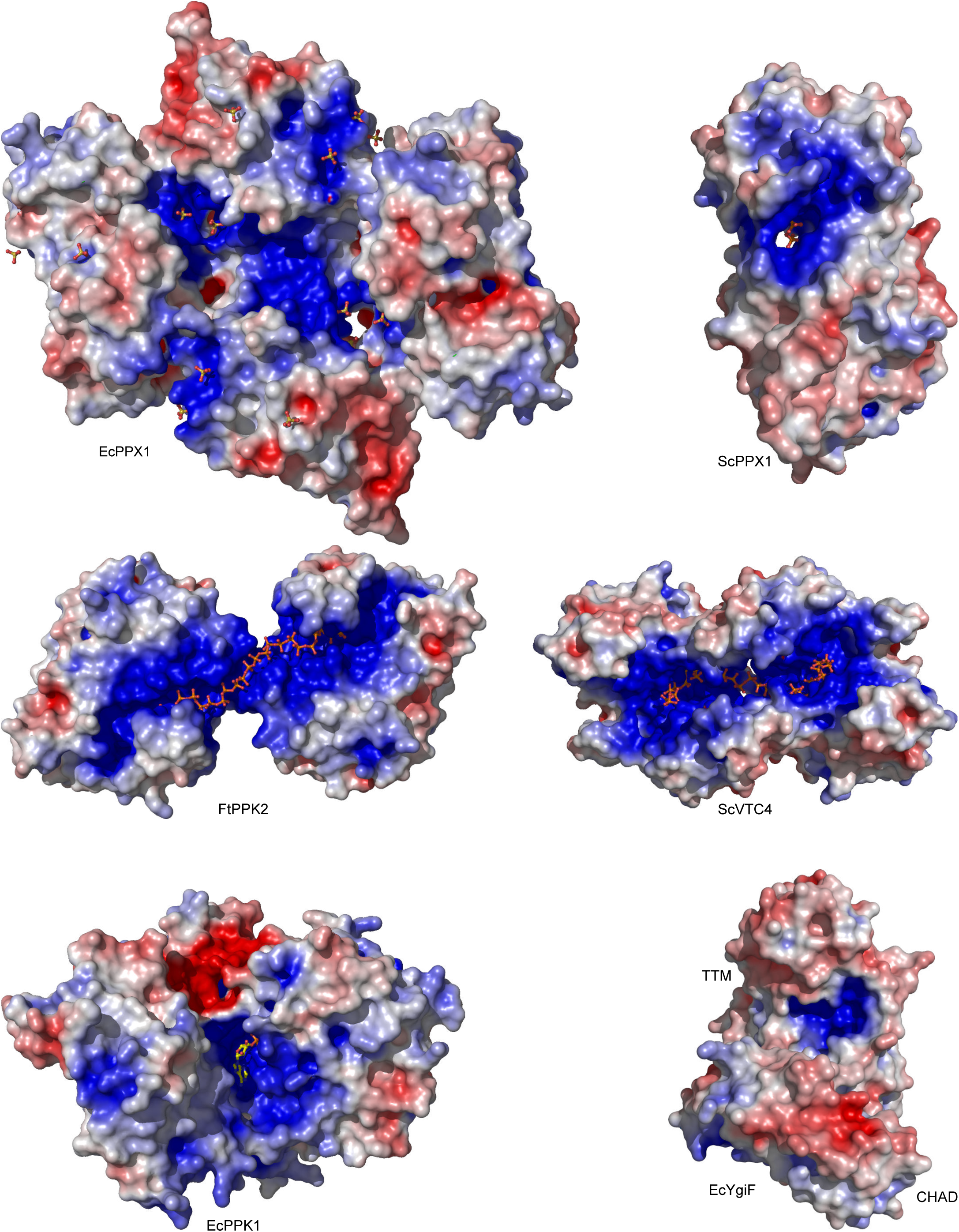
PolyP-metabolizing enzymes contain large basic surface patches facilitating polyP binding. Depicted are electrostatic potentials calculated in APBS (65), mapped onto molecular surfaces calculated in Pymol for EcPPX1 (PDB-ID: 1u6z), ScPPX1 (PDB-ID: 2qb7), FtPPK2 (PDB-ID: 5llf), ScVTC4 (PDB-ID: 3g3q), EcPPK1 (PDB-ID: 1xdp) and ygiF (PDB-ID: 5a61). Bound phosphates, sulfates, polyPs and PPK1 substrate analogs are highlighted in bonds representation.

**SI Appendix Fig. S6.**
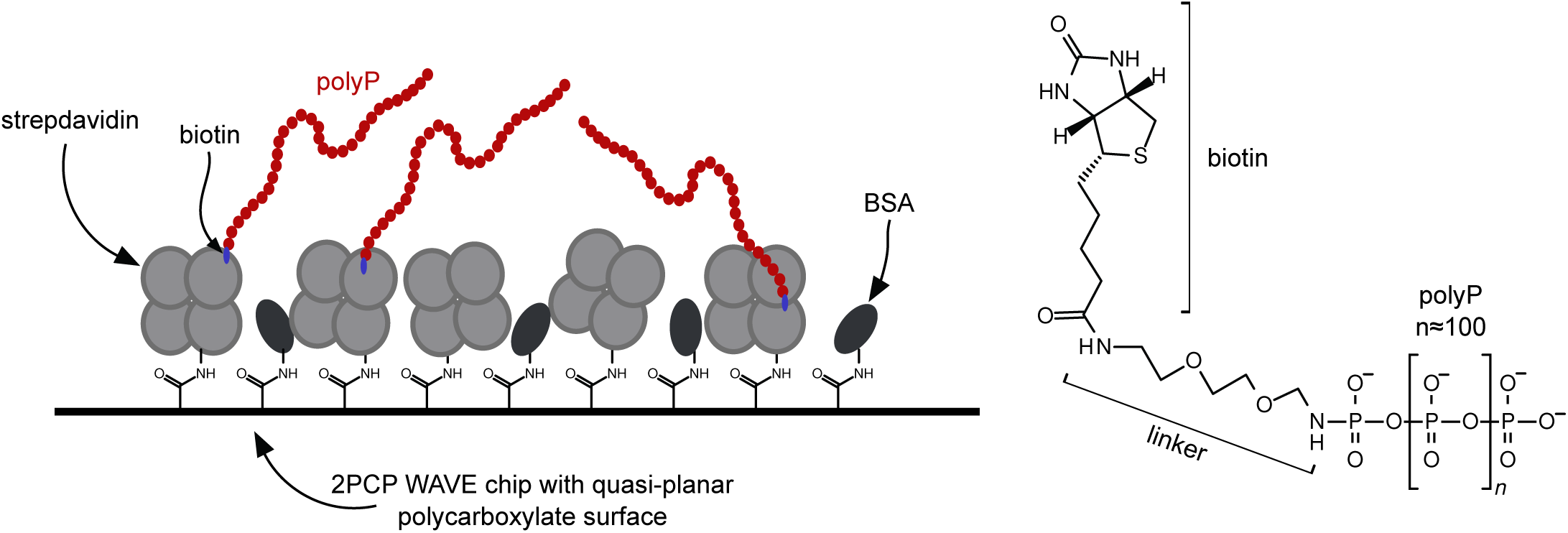
Schematic overview of the GCI chip used to determine the interaction between CHAD domain-containing proteins and polyP. Streptavidin was covalently linked to a 4PCP-WAVE chip (Creoptix AG), and subsequently passivated with BSA (left). Biotinylated polyP (average chain-length ∼ 100 P_i_ units, right) was then bound to the immobilized strepdavidin on the GCI chip.

**SI Appendix Fig. S7.**
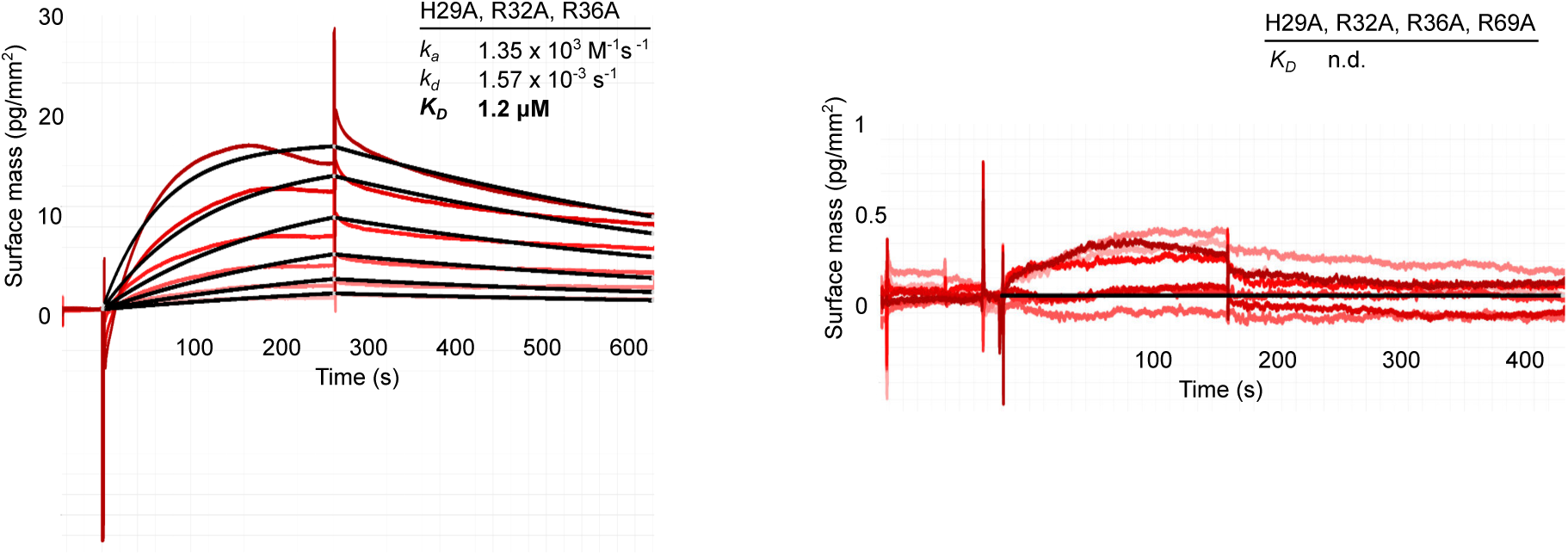
Mutations of conserved basic amino-acids in SsCHAD disrupt polyP binding in quantitative GCI assays. Shown are sensograms (in red), the respective fits (in black), and table summaries of the derived association rate constant (k _*a*_), dissociation rate constant (k_*d*_) and dissociation constant (K_*D*_). Simultaneous mutation of His29 (His253 in CtCHAD), Arg32 (Arg256 in CtCHAD) and Arg36 (Arg260 in CtCHAD) to alanine strongly reduces binding of the mutant protein to polyP (wild-type K_*D*_ is 44.5 nM). Additional mutation of Arg69 to Ala (Arg296 in CtCHAD) disrupts binding in our GCI assay.

**SI Appendix Fig. S8.**
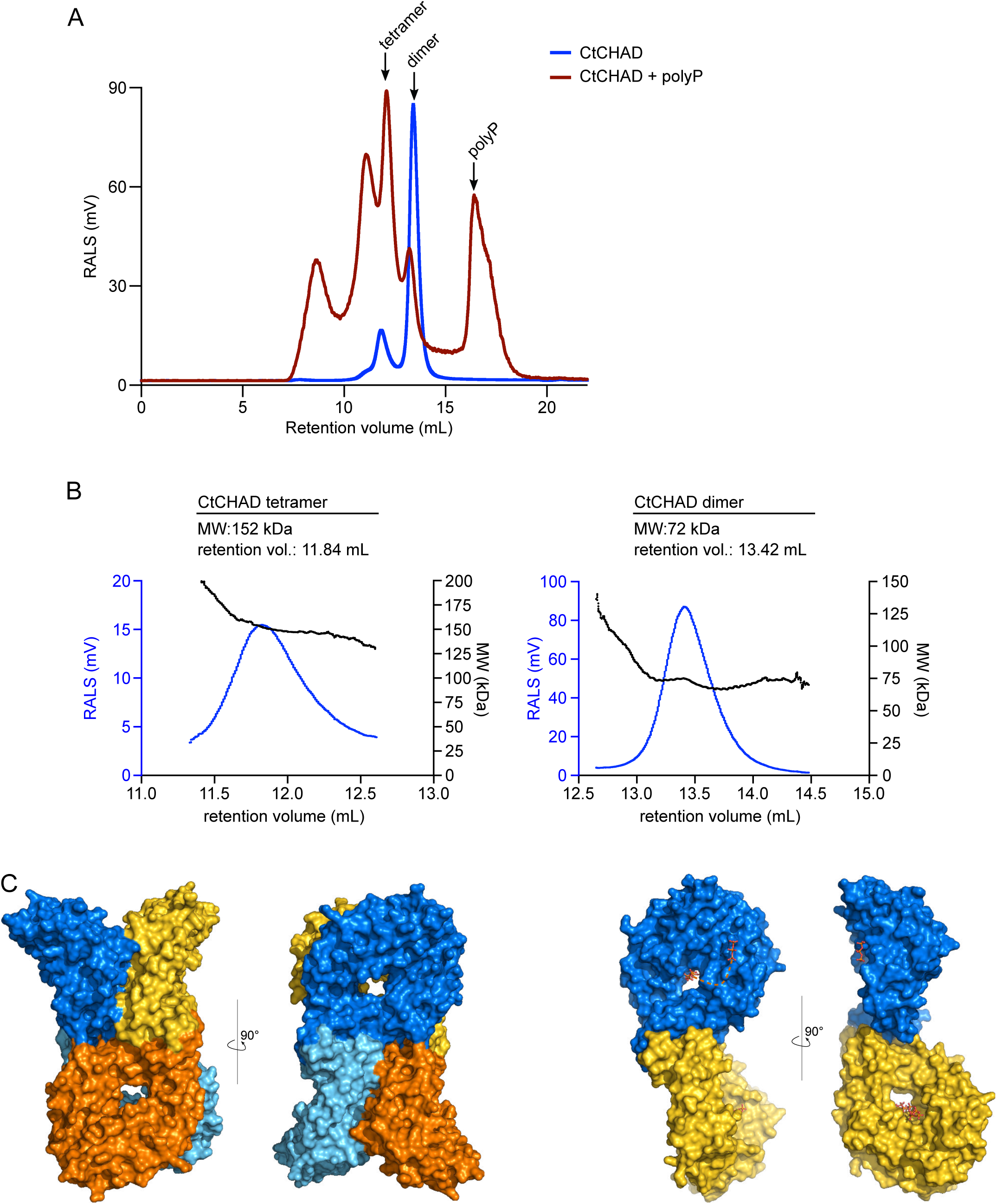
PolyP-binding may induce oligomerisation of CtCHAD. Size-exclusion chromatography (SEC) and right angle light scattering (RALS) analysis of CtCHAD in complex with polyP. **A**, RALS traces of CtCHAD in the presence (red) and absence (blue) of 1 mM polyP (average chain length ∼ 30 P_i_ units). Indicated oligomeric states are calculated based on CtCHAD, as shown in panel B. **B**, RALS traces (blue) and extrapolated molecular weight (black) of the two peaks of CtCHAD. Retention volumes and calculated molecular weights are shown alongside. In the absence of polyP, CtCHAD forms a dimer (∼ 90 %) and a tetramer (∼ 10 %). The theoretical molecular weight of monomeric CtCHAD is 36,194 Da. C, Putative tetrameric and dimeric assemblies for CtCHAD based on crystal packing analysis as implemented in the program PISA (62). Shown are molecular surfaces of a CtCHAD tetramer (left) and dimer (right) in two orientations, respectively. Note that the position of the central cavities in the tetramer and the polyP binding surfaces in the dimer could allow for the cooperative binding of long-chain polyPs.

**SI Appendix Table S1.** Crystallographic data collection and refinement.

|  | RcCHAD | CtCHAD – apo (sulfate) | CtCHAD – polyP |
| --- | --- | --- | --- |
| **PDB-ID** | **6QV5** | **6QV7** | **6QVA** |
| **Data collection** |  |  |  |
| Space group | *P* 6<sub>1</sub> 2 2 | *I* 2 2 2 | *I* 2 2 2 |
| Cell dimensions |  |  |  |
| *a*, *b*, *c* (Å) | 116.4, 116.4, 119.3 | 69.5, 95.7, 102.8 | 69.4, 97.0, 106.4 |
| α, β, γ (°) | 90, 90, 120 | 90, 90, 90 | 90, 90, 90 |
| Resolution (Å) | 41.65 – 2.30 (2.44 – 2.30) | 36.80 – 1.72 (1.82 – 1.72) | 19.70 – 2.05 |
| *R*<sub>meas</sub><sup>#</sup> | 0.172 (3.68) | 0.074 (2.21) | 0.155 (2.35) |
| CC(1/2)<sup>#</sup> | 0.99 (0.50) | 1.0 (0.50) | 1.0 (0.43) |
| *I*/σ *I*<sup>#</sup> | 15.5 (1.0) | 16.0 (1.0) | 14.1 (1.1) |
| Completeness (%)<sup>#</sup> | 99.9 (99.6) | 100 (99.8) | 99.6 (99.6) |
| Redundancy<sup>#</sup> | 21.2 (21.5) | 7.0 (7.1) | 13.3 (13.2) |
| Wilson B-factor<sup>#</sup> | 62.5 | 39.3 | 46.1 |
| **Refinement** |  |  |  |
| Resolution (Å) | 41.65 – 2.30 | 36.8 – 1.72 | 19.70 – 2.05 |
| No. reflections | 21,737 | 70,305 | 21,397 |
| *R*<sub>work</sub>/ *R*<sub>free</sub><sup>\$</sup> | 0.232 / 0.242 | 0.189 / 0.222 | 0.208 / 0.255 |
| No. atoms |  |  |  |
| protein | 2,237 | 2,584 | 2,518 |
| ligand | 5 | 88 | 71 |
| solvent | 27 | 179 | 68 |
| Res. B-factors<sup>\$</sup> | | | |
| protein | 84.2 | 42.6 | 49.7 |
| ligand | 74.8 | 71.6 | 98.2 |
| solvent | 56.0 | 51.4 | 49.5 |
| R.m.s deviations<sup>\$</sup> | | | |
| bond lengths (Å) | 0.011 | 0.01 | 0.01 |
| bond angles (°) | 1.49 | 1.07 | 1.73 |
| Ramachandran plot<sup>\$</sup>: | | | |
| most favored regions (%) | 98.25 | 99.36 | 98.70 |
| outliers (%) | 0 | 0 | 0 |
| MolProbity score<sup>\$</sup> | 1.25 | 1.24 | 1.01 |

Highest resolution shell is shown in parenthesis.
<sup>#</sup>as defined in XDS (63)
<sup>\*</sup>as defined in autoBUSTER (Global Phasing Limited) (RcCHAD), phenix.refine (54) (CtCHAD apo) or Refmac5 (58) (CtCHAD - polyP)
<sup>\$</sup>as defined in Molprobity (64)

